# New Techniques for Ancient Proteins: Direct Coupling Analysis Applied on Proteins involved in Iron Sulfur Cluster Biogenesis

**DOI:** 10.1101/103283

**Authors:** Marco Fantini, Duccio Malinverni, Paolo De Los Rios, Annalisa Pastore

**Affiliations:** Scuola normale superiore (SNS), Pisa, Italy; Laboratoire de Biophysique Statistique, Faculté de Sciences de Base, École Polytechnique Fédérale de Lausanne - EPFL, 1015 Lausanne, Switzerland; Institute of Bioengineering, School of Life Sciences, École Polytechnique Fédérale de Lausanne - EPFL, 1015 Lausanne, Switzerland; Maurice Wohl Institute, King’s College, London, UK; Molecular Medicine Department, University of Pavia, Pavia, Italy

## Abstract

Direct coupling analysis (DCA) is a powerful tool based on protein evolution and introduced to predict protein fold and protein-protein interactions which has been applied also to the prediction of entire interactomes. We have used DCA to analyse three proteins of the iron-sulfur biogenesis machine, an essential metabolic pathway conserved in all organisms. We show that, although based on a relatively small number of sequences due to its distribution in genomes, we can correctly recapitulate all the features of the fold of the CyaY/frataxin family, a protein involved in the human disease Friedreich’s ataxia. This result gave us confidence in the use of this tool. Application of DCA to the iron-sulfur cluster scaffold protein IscU, which has been suggested to function both as an ordered and a disordered form, allows us to clearly distinguish evolutionary traces of the structured species, suggesting that, if present in the cell, the disordered form has not left any evolutionary imprinting. We observe instead, for the first time, direct indications of how the protein can dimerize head-to-head and bind 4Fe4S clusters. Analysis of the alternative scaffold protein IscA provides strong support to a coordination of the cluster mediated by a dimeric rather than a tetrameric form as previously suggested. Our analysis also suggests the presence in solution of a mixture of monomeric and dimeric species and guide us to the prevalent one. Finally, we used DCA to analyse protein-protein interactions between some of these proteins and discuss the potentialities and the limitations of the method.

## I. INTRODUCTION

The whole history of protein folding and interactions is encoded in the correlations between residues in the protein sequence. The logic connecting residue-residue contacts to evolutionary correlation is very simple: residues in contact cannot evolve independently. If one residue gets larger, the other needs to be smaller in a concerted and not necessarily pairwise way. Charges must be compensated in the same way. Stabilizing/destabilizing amino acid substitutions need to be compensated by substitution of other interacting positions over the evolutionary timescale to retain interaction. In principle, one could use a comparative analysis of the primary sequences of proteins as a powerful way to predict their structures and interactions. This idea has been the “elusive Holy Grail” for more than twenty years since the first establishment of bioinformatics [1]. More recently an effective method, named direct coupling analysis (DCA) [2,3], has been proposed as a powerful approach to determine which residues interact the most from an evolutionary perspective, exploiting the large, and growing, number of available protein sequences. The method has been successfully used to acquire constraints for structural, dynamical and functional analysis [4-7], multimerization [8,9], and to shed light on interaction specificity [10] and inter-pathway cross-talk in bacterial signal transduction [11].

Here, we have applied DCA to explore the nature of the interactions between proteins involved in the iron-sulfur (FeS) cluster biogenesis pathway. Iron-sulfur clusters are essential prosthetic groups in biological material bound to proteins to provide electrons in reduction/oxidation reactions and/or stabilize protein folds. Their biosynthesis is a complex process involving specialized machines which mediate the recruitment of sulfur and free iron from the cellular environment, catalyse the synthesis and fulfil the delivery of the newly formed clusters to acceptor proteins. In bacteria, the systems able to perform these tasks belong to the nif (nitrogen fixation, *Nif*iscA-nifSU), isc (iron-sulfur complex, *isc*RSUA-*hsc*BA-*fdx*) and suf (mobilization of sulfur, *suf*ABCDSE) operons. Amongst these, the most universal one is the isc operon, whose proteins have direct orthologues in eukaryotes. Because malfunction in FeS cluster assembly has direct effects onto human health [12,13], elucidating the structures and interaction patterns between the various proteins involved in this process can provide valuable insights in the origin of several diseases.

The central players in the isc machinery are IscS (or Nfs1 in eukaryotes) and IscU (Isu). IscS is a desulfurase, which converts cysteine to alanine and forms the persulfide that participates to the cluster, and IscU is a scaffold protein where the cluster is assembled. Together, they form a complex in which two IscU monomers are bound to the IscS obligate dimer. IscU was suggested to exist in the cell in two conformational states, one folded and ordered (S state), the second being partially unfolded (D state) [14]. However, all crystal structures of IscU in isolation and in complexes with zinc or IscS capture the protein in its ordered state. Two regulatory proteins are CyaY (frataxin), which is the protein involved in Friedreich’s ataxia in humans, and IscA thought to be an alternative scaffold protein. CyaY/frataxin is a monomeric protein formed by a globular conserved domain, in eukaryotes preceded by an intrinsically unfolded mitochondrial import sequence. It is highly conserved from bacteria to primates [15], to act as a regulator of the enzymatic activity of IscS and to bind it in a site close to the enzyme active site [16,17]. Puzzlingly, its presence seems to inhibit the activity in prokaryotes but to activate it in eukaryotes [18-21]. IscA is an ancient protein thought to be an alternative scaffold for cluster formation. The IscA family is characterized by a conserved CXnCGCG pattern though to be involved in iron and/or 2Fe-2S binding [22]. In all available structures, IscA is either dimeric or tetrameric but different symmetries and cluster coordination were suggested. We used DCA to address important outstanding questions, which would help us to understand the specific role of these proteins and their fold. We used frataxin, which is monomeric and globular, to calibrate the method. We then tested whether any trace of the D state of IscU is detectable as compared to the S state and whether evolution provides information on the quaternary arrangement of IscA. We found that this technique was able to describe in great detail the proteins considered. We were able to identify the correct biological location of the elusive N-terminus of the IscU protein and conserved contacts which hint at a head-to-head dimerization of the protein which is in agreement with the cluster coordination. We also found that not all the IscA structures in the PDB database match the conserved contacts which suggests that the location of the FeS cluster was likely misattributed. Finally, we used DCA to predict protein interactions. We could predict successfully interactions between IscU and the functional partner IscS whereas contacts predicted for CyaY do not match our current knowledge. These observations are likely to reflect the possibilities but also the limitations of DCA.

## II. RESULTS

### A. Validating the method on the frataxin family

The major sequence divergence within the CyaY/frataxin family is in the non-conserved mainly unstructured N-terminus [23,24]. The evolutionary conserved C-terminal domain forms a compact globular structure in which two *α*-helices pack against *β*-sheet composed of 5-7 strands arranged in a *αβββββ*(*ββ*)*α* motif. The available structures of this region are all similar (average RMSD 2.3 Å) with minor differences in details (Table 1). Different orthologs differ for the length of the C-terminus that is longer in human frataxin and shorter in yeast. This difference contributes to the thermodynamic stability of the protein [25]. Experimental evidence suggests that the region interacting with iron and with the desulfurase IscS/Nfs1 is located in *α*1 and *β*1 (Fig 1A) [18,26,27].

We retrieved from the Uniprot database all the sequences matching a Hidden Markov Model (HMM) constructed from a seed made of the 196 CyaY entries of the Swiss-Prot database. We then built a multiple sequence alignment (MSA) containing 3459 sequences, defining 109 consensus residue positions which cover 1102 eukaryotes and 2326 bacteria. The number of retrieved sequences is relatively small for a successful application of DCA but reflects the absence of frataxin in several species [28]. We then performed DCA on this MSA using the pseudo-likelihood approximation as described in Balakrishnan et al. [29], a method that estimates the joint probability distribution of a collection of random variables. The predicted contacts are displayed in contact maps which have the protein sequence numbering on both axes. Contacts are displayed as spots which indicate interactions between residues. Traces antiparallel to the diagonal indicate that this region forms antiparallel secondary structure. Parallel traces reflect interactions between parallel strands. Contacts which do not line up in parallel or antiparallel fashion but cluster in various regions of the plot correspond to contacts between distal elements.

The predicted contacts (Fig 1B) were ranked according to their DCA scores, which describe the coevolution strength between pairs of residues. Comparison of the frataxin structures with the contacts predicted by DCA, discarding contacts between residues less than five residues apart which reflect local secondary structures, shows a perfect match for the first 25 residues, drops to two thirds in the first hundred, and stays over 50% at the 250 residue mark (Fig 1C). We retained the top 109 DCA contacts with the highest scores which correspond to 2% of the total 5460 possible contacts. The retained contacts correlate well with the secondary structure of the protein. Additionally, three clusters were observed, all involving the domain N-terminus (Fig 1D). They reflect packing of *α*1 against *β*1-*β*2 and *β*3-*β*4. The third cluster reflects the contacts between the two helices. This tells us how important *α*1 is for this protein fold. The only other tertiary interactions between distant secondary structure elements involve *β*4-*β*5 and the C-terminal *α*2. This interaction is reflected in the DCA analysis by a small cluster visible at the very bottom of the DCA plot.

These results support the confident use of DCA for the analysis of FeS proteins: even though the number of retrieved sequences is suboptimal, we could recapture all the important features of the CyaY/frataxin fold.

All the crystal structures of IscU have a folded N-terminus while most of the NMR structures show an unstructured or partially structured N-terminus. Only a few structures are available for IscA, all from X-ray crystallography.

**TABLE I.** Summary of the available CyaY/frataxin structures

| PDB ID | ORIGIN | TYPE | LIGANDS | MUTATIONS | DISPERSION | RELEASE DATE |
| --- | --- | --- | --- | --- | --- | --- |
| 1EW4 | E.coli (B) | X-RAY | - | - | Monomer | Aug2000 |
| 1SOY | E.coli (B) | NMR | - | - | Monomer | Nov 2004 |
| 2EFF | E.coli (B) | X-RAY | Co<sup>2+</sup> | - | Monomer | Oct 2007 |
| 2P1X | E.coli (B) | X-RAY | Eu<sup>3+</sup> | - | Monomer | Oct 2007 |
| 4HS5 | P.ingrahamii (B) | X-RAY | - | - | Dimer | Mar 2013 |
| 4LK8 | P.ingrahamii (B) | X-RAY | Co<sup>2+</sup> | - | Dimer | Jul 2014 |
| 4LP1 | P.ingrahamii (B) | X-RAY | Eu<sup>3+</sup> | - | Dimer | Jul 2014 |
| 4JPD | B.cenocepacia (B) | X-RAY | - | - | Monomer | Mar 2013 |
| 4EC2 | S.cerevisiae (YM) | X-RAY | Fe<sup>3+</sup> | res52-174,Y73A | Trimer | Jan 2013 |
| 3OEQ | S.cerevisiae (YM) | X-RAY | - | res52-174,Y73A | Trimer | Aug 2011 |
| 3OER | S.cerevisiae (YM) | X-RAY | Co<sup>2+</sup> | res52-174,Y73A | Trimer | Aug 2011 |
| 2FQL | S.cerevisiae (YM) | X-RAY | - | res52-174,Y73A | Trimer | Nov 2003 |
| 2GA5 | S.cerevisiae (YM) | NMR | - | - | Monomer | Mar 2006 |
| 1EKG | H.sapiens (HM) | X-RAY | - | - | Monomer | Nov 2000 |
| 1LY7 | H.sapiens (HM) | NMR | - | res91-210(C-term) | Monomer | Jun 2002 |
| 3S4M | H.sapiens (HM) | X-RAY | - | - | Monomer | Jun 2011 |
| 3S5D | H.sapiens (HM) | X-RAY | - | W155A | Monomer | Jun 2011 |
| 3S5E | H.sapiens (HM) | X-RAY | - | W155R | Monomer | Jun 2011 |
| 3S5F | H.sapiens (HM) | X-RAY | - | W155F | Monomer | Jun 2011 |
| 3T3J | H.sapiens (HM) | X-RAY | - | N146K | Monomer | Aug 2011 |
| 3T3K | H.sapiens (HM) | X-RAY | - | Q148R | Monomer | Aug 2011 |
| 3T3L | H.sapiens (HM) | X-RAY | - | Q153A | Monomer | Aug 2011 |
| 3T3T | H.sapiens (HM) | X-RAY | - | Q148G | Monomer | Aug 2011 |
| 3T3X | H.sapiens (HM) | X-RAY | - | R165C | Monomer | Aug 2011 |

**TABLE II.** Summary of the available IscU structures.

| PDB | STRUCTURED N-TERMINUS<sup>a</sup> | TYPE | LIGANDS | MUTATIONS | DISPERSION | RELEASE DATE |
| --- | --- | --- | --- | --- | --- | --- |
| 2Z7E | ✓ | X-RAY | 2Fe-2S | D38iscUA | Homotrimer | Aug 2008 |
| 4EB7 | ✓ | X-RAY | 2Fe-2S, PLP | V16iscSI | IscU-(IscS2) trimer | May 2012 |
| 4EB5 | ✓ | X-RAY | 2Fe-2S, PLP | - | (IscU-IscS)2 tetramer | May 2012 |
| 3LVL | ✓ | X-RAY | PLP | - | IscU-IscS dimer | Apr 2010 |
| 1SU0 | ✓ | X-RAY | Zn<sup>2+</sup> | - | Monomer | Aug 2004 |
| 2QQ4 | ✓ | X-RAY | Zn<sup>2+</sup> | - | Homodecamer | Jul 2008 |
| 1R9P | ✗ | NMR | Zn<sup>2+</sup> | - | Monomer | Nov 2004 |
| 1Q48 | ✗ | NMR | - | - | Monomer | Nov 2003 |
| 2L4X | (✗) | NMR | - | - | Monomer | Dec 2011 |
| 2KQK | (✗) | NMR | - | D39iscUA | Monomer | Nov 2010 |
| 1XJS | ✓ | NMR | Zn<sup>2+</sup> | - | Monomer | Jan 2005 |
| 1WFZ | (✓) | NMR | Zn<sup>2+</sup> | - | Monomer | Nov 2004 |

<sup>a</sup> N-terminus overlapping with the predicted DCA contacts. In bracket partial/incomplete overlap.

### B. Structure of IscU proteins and N-terminal localization

IscU is a more complex case. Twelve structures are available from 8 different species (Table 2). They can be divided in three groups. All the X-ray structures, which are available for isolated cluster-loaded (holo), zinc-loaded and cluster-free (apo) IscU as well as complex with IscS/Nfs1, have a compact ordered structure with a *β*-sheet packing against two *α*-helices (Fig 2A). The N-terminus (residues 1-21) does not contain regular secondary structure elements except for a two-turn helix (*α*1) between residues 5-12 which packs against the other helix anchoring the N-terminus to the rest of the structure. In one of the structures (2Z7E), the N-terminus adopts different orientations in the different protomers of a homo-trimer. In the solution structures, (1R9P, 1Q48, 2L4X, 2KQK and the 1WFZ), the fold is similar but the N-terminus is disordered and completely solvent-exposed (Fig 2B). Some of these structures are thought to contain a zinc atom in the same position where the cluster is coordinated (i.e. on the tip of the approximately ellipsoid where three conserved cysteines are). However, zinc is NMR silent and could not be observed directly. Only two crystallographic structures (1SU0 and 2QQ4) explicitly contain zinc. Finally, one zinc free NMR structure (2L4X) is supposed to be representative of the D state.

**TABLE III.** Summary of the available IscA and IscA-like structures.

| PDB | PROTEIN | TYPE | LIGANDS | MUTATIONS | DISPERSION | RELEASE DATE |
| --- | --- | --- | --- | --- | --- | --- |
| 1R94 | E.coli IscA | X-RAY | Hg<sup>2+</sup> | - | Dimer<sup>a</sup> | Dec 2005 |
| 1R95 | E.coli IscA | X-RAY | - | - | Dimer<sup>a</sup> | Dec 2005 |
| 1S98 | E.coli IscA | X-RAY | - | L69M | Dimer<sup>a</sup> | Jun 2004 |
| 1X0G | T.elongatus IscA | X-RAY | 2Fe-2S | - | Tetramer | Jun 2006 |
| 2D2A | E.coli SufA | X-RAY | - | - | Dimer | Dec 2005 |
| 2K4Z | A.vinosum DsrR | NMR | - | - | Monomer | Jul 2008 |

<sup>a</sup> The authors propose the tetramer as the functional unit.

**FIG. 1.**
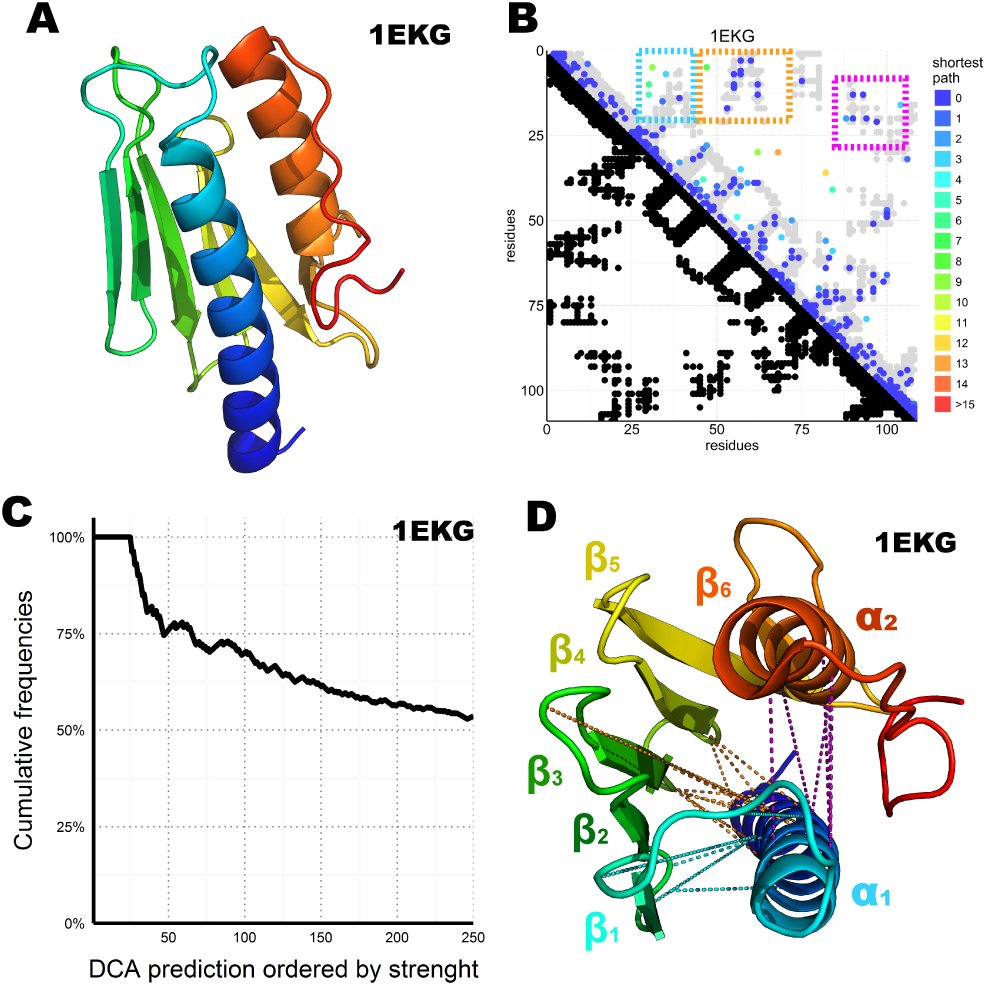
DCA prediction of interacting residues in the CyaY/frataxin family. A) CyaY/frataxin reference structure 1EKG. B) The residue number of the family consensus sequence from the N- to the C-terminus is displayed on both axes. In the bottom half of the plot, black dots are used to indicate residues in contact in the 1EKG reference structure. In the top half of the plot, the predicted DCA contacts are coloured by the shortest path from a reference matching position (gray dots). The gray dots are the same shown in black in the bottom half but plotted again to help visualisation. The tree major clusters are highlighted by coloured frames. C) Accuracy of the DCA as a function of the number of considered top scoring residues. The plot shows the frequency of reference-matching predictions (number of matching hits divided by the number of prediction considered up to that point) over the DCA contacts sorted by strength. D) 1EKG structure with the three main DCA predicted cluster contacts. In cyan: the *α*1-*β*1*β*2 cluster; in orange: the *α*1-*β*3*β*4 cluster; in purple: the *α*1-*α*2 cluster.

It is distorted and contains only a *β*-hairpin and the C-terminal helix. It is probably more correct to describe this entry as a nascent chain or a molten globule rather than a structure as we normally intend it. Its presence in PDB is misleading.

DCA analysis on 13148 IscU sequences, resulted in clear coevolutionary prediction of contacts (Fig 2C,D). Using the secondary structure and the nomenclature described in the IscU alignment [30], we can observe interactions between secondary structure elements: the contacts between *β*1-*β*2, *β*2-*β*3, *β*3-*α*2, *α*2-*α*3, and *α*3-*α*6 left traces perpendicular to the diagonal, while the *β*2-*α*2, *β*3-*α*6, *β*2-*α*6 interactions are reflected by three parallel traces. All secondary structure elements between *β*1 and *α*6 form contacts with the previous and the subsequent secondary elements, forming hairpins. The parallel traces reflect interactions between parallel strands. The *α*1 helix is excluded from this pattern and forms interactions with several strands suggesting a transversal orientation which crosses the sheet.

Most experimental structures agree with these predicted contacts (Fig 2C,D) with the exception of the N-terminal region (up to ca. residue 16) which is also where the structures differ most. Contacts between the N-terminus and the *β*2-*β*3-*α*2 region are conserved, in support to a structured state of the *α*1 region (Fig 2E,F). This does not, however, preclude the existence or the functional relevance of a disordered conformation of the N-terminus: disordered regions would likely not have a co-evolutionary signal and are thus out of reach in current DCA predictions.

The N-terminus also forms contacts with the *β*-sheets and the *α*1-*β*1 loop. Superposition of the predicted contacts to the deposited structures leaves two unaccounted predicted contact clusters, one between *α*2 and *β*1-*β*2 loop, another within the *α*5 region (Fig 3A). These contacts are incompatible with the inter-molecular interactions observed in the crystal structures of the cluster-loaded trimer (2Z7E) or of a decamer (2QQ4) (Fig S1) and include areas involved in or surrounding the FeS cluster binding site (Fig 3B). A different explanation could be that these contacts reflect formation of a head-to-head dimer with an interface located around the conserved cysteines. This hypothesis would be fully consistent with the necessity of at least a dimer to coordinate a 4Fe4S cluster [31] according to a oxidative mechanism previously proposed [32].

**FIG. 2.**
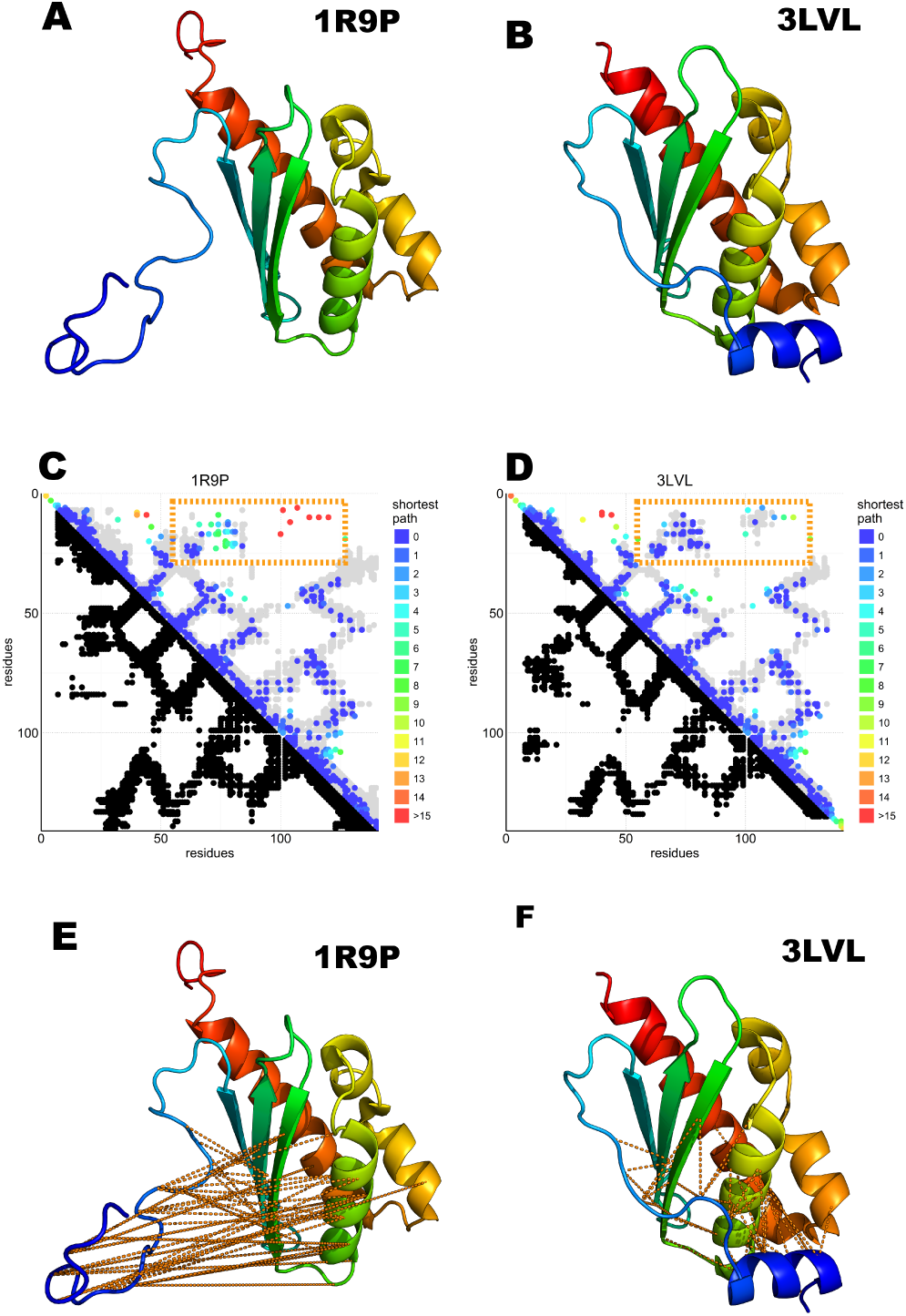
Predicted N-terminal interactions over IscU models with structured (3LVL) or unstructured (1R9P) N-terminus. A) IscU reference structure 3LVL illustrating the structured N-terminus. B) IscU reference structure 1R9P, prototypical of the NMR structures with an unstructured N-terminus. C-D) DCA on the IscU family over IscU models with structured (3LVL) or unstructured (1R9P) N-terminus. Each axis contains the family consensus sequence from the N-to the C-terminus. Orange frames highlight the contacts missing in the unstructured (A) but present in the structured (B) N-terminus. E-F) The missing contacts are compared to the structures with ordered (3LVL) and disordered (1R9P) N-terminus.

**FIG. 3.**
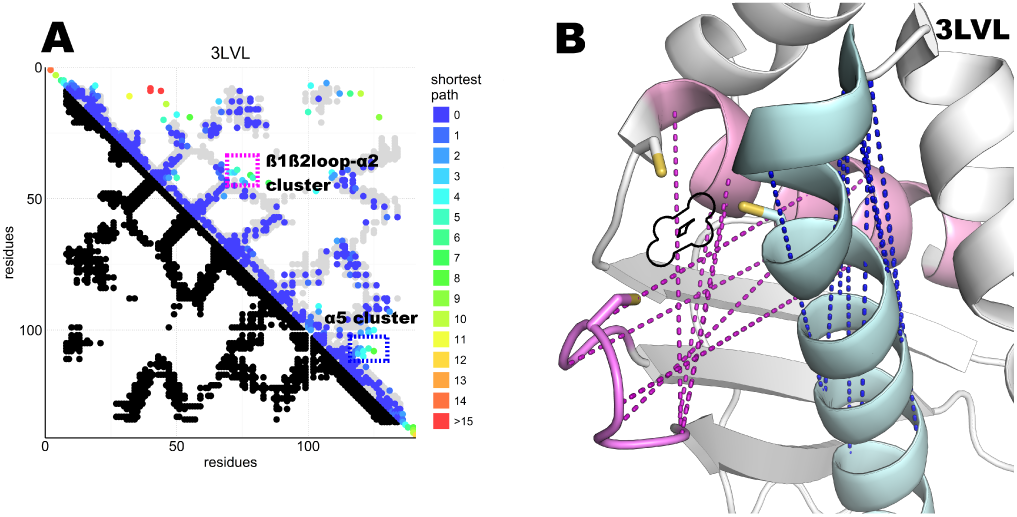
DCA Contact map of the IscU protein and structure showing unaccounted contact clusters. A) DCA on the IscU family compared to the 3LVL structure. The two unaccounted clusters are highlighted by coloured frames. B) FeS cluster binding site of IscU (3LVL) with the unaccounted contacts between the *β*1-*β*2 loop and α2 (pink) and within *α*5 (cyan). In purple, the unaccounted loop-helix predicted contacts; in blue, the *α*5 cluster. A black outline indicates the location of the FeS cluster in the structure. All the involved residues are at or close to the active site. Cysteines side chains are shown explicitly.

### C. Multimerization and FeS cluster coordination of IscA

Seven structures of IscA-like proteins are available (Table 3). The first published structure (1R95) [33] has an internal two-fold symmetry with tandem pseudo-symmetric motifs (*β*1-*α*1-*β*2-*β*3/*β*5-*α*2-*β*6-*β*7) separated by a quasi-palindromic hinge (E_43_FVDEPTPEDIVFE_56_ in the *β*3-*α*4 region). The fold of each protomer consists of a *β*-sandwich of a mixed twisted four-stranded *β*-sheet, *β*4-*β*5-*β*2-*β*3, packed against a three-stranded *β*1-*β*6-*β*7 sheet. The protomers could form a dimer or two possible tetramers or dimer of dimers (tetramers A and B, Fig 4A). The electron density around the C-terminus (where two of the three cysteine residues are) is fuzzy, indicating disorder or conformational exchange. An alternative apo IscA crystal structure [34] has the individual protomers nearly identical to those observed in 1R95 but the dimer interface, described as an *α*1*α*2 dimer with minor differences between protomers, is different. The overall tetrameric (*α*1*α*2)2 structure is similar to the 1R95 A tetramer. Also this structure lacks a defined C-terminus but the authors modelled it based on stereochemical parameters. The authors concluded that the cysteines of the dimer would be unable to coordinate the FeS cluster and that tetramer formation is necessary to stabilise coordination. They also suggested that of the three cysteines of the C*X*_*n*_CGCG motif, only the last two (Cys99 and Cys101 in E. coli) are involved in cluster coordination, whereas Cys35 would remain idle. The only fully resolved holo IscA is from *T. elongatus* (1X0G). This structure has a structured C-terminus which allows coordination of the FeS cluster. It is a dimer of asymmetric dimers (*αβ*)2 and has domain swapping between two of the protomers (*β* and *β*’) which exchange their central domain forming a long intertwined *β*-sheet (Fig 4B). The unusual asymmetry imposes asymmetric interfaces, one of which (the one between *α*and the domain-swapped *β*’) forms the pocket which accommodates the FeS cluster. The pocket itself is asymmetric with the cysteine motif (Cys37, Cys101, Cys103) contributed both by the *α*protomer and the swapped *β*domain of the protomer (Cys103 (β_*sw*_) (Fig 4B and Fig S2).

Most of the sequences belong either to the IscA or to the ErpA subfamilies but comprise also SufA and the eukaryotic paralogs IscA1/IscA2 (ca. 11,000 sequences). These proteins are all part of the A-type carrier (ATC) family and should have overlapping functions. Structurally, both SufA (2D2A) and IscA (1R95, 1S98) have similar contact maps except for two regions, which account for contacts within the C-terminus and between the C-terminus and residues 30-40. These regions contain the three conserved cysteines. Since cluster coordination is thought to occur inter-molecularly because no structure could allow intra-molecular coordination [35], we hypothesize that these contacts reflect inter-molecular interactions. None of the inter-chain contact maps matches sh he experimental structures (data not shown), strongly suggesting that in solution there might be different structures in mutual equilibrium or that none of the available structure represents the functional species. The first hypothesis is also in agreement with the diversity of packing observed in the crystal structures.

The contacts within C-terminal residues show the characteristic pattern of *β*-sheets or loop conformations. These patterns could be in agreement with the anomalous swapped dimer of 1X0G, where the loop harbouring the first cysteine of the CX_*n*_CGCG motif (Cys37) is bent towards the C-terminus and stabilized by steric hindrance of the swapped central twisted *β*-sheets. In this structure, cluster coordination is asymmetric and achieved by Cys37 and Cys101 of the *α* protomer and Cys103 of the *β* protomer. The evolutionary trace of contacts between the C-terminus (residues 98-112) and the loop between residues 33-41 suggests the existence of a conformation which allows the proximity of the first cysteine (Cys37) to the terminal cysteine pair (Cys101 and Cys103) (Fig S2), supporting a contribution of Cys37 in cluster coordination. This conclusion is at strong variance with the previous belief that only the C-terminal cysteines participate to coordination and implies that cluster coordination can occur at the level of the dimer without invoking formation of a tetramer. The 1X0G structure is currently the only available structure able to describe cluster coordination although domain swapping may not be required to explain the interactions: domain swapping could easily be replaced by a non-swapped protomer in a symmetric dimer.

We can thus conclude that DCA analysis of IscA suggests new important hypotheses which can change drastically our views on this protein cluster coordination properties.

### D. Protein-protein interactions

DCA can in principle be extended to predict conserved contacts between interacting proteins on the basis of MSAs of protein pair sequences that are known to interact. In the absence of such a curated set, several matching strategies have been developed [8–10,36]. Among these, two independent implementations have recently been suggested in back-to-back publications [10,36]. We adopted the Iterative Paralog Matching (IPA) [10] to investigate the interactions between frataxin, IscU and the desulfurase IscS and used a self-consistent method that simultanously identifies the best matching sequences among paralogs, and predicts pairs of interacting residues across two proteins. In this approach, multiple IPA runs are performed, and protein-protein contacts are scored based on the number of times they are accepted among all the runs (acceptance frequency). We first analyzed the interactions between IscU and IscS, because a high resolution crystal structure of this complex is available (3LVL). We observed that the four most often accepted contacts do indeed lie in the interface of the IscU-IscS dimer. These contacts have acceptance frequencies between 100% and 85% (Fig 5A and Fig S3). Contacts with lower acceptance frequencies are mainly incompatible with the structural model of the IscU-IscS dimer (i.e. false-positives). We also observed at least one contact (V17-L383, accepted in 17% IPA runs), that lies in the IscU-IscS interface. In the absence of an absolute scale quantifiying the reliability of predicted contacts, and of known structures for the IscU-frataxin and IscS-frataxin complexes, we used the IscU-IscS case as a reference. We assumed that contacts being accepted in more than 85% of IPA simulations are all in excellent agreement with an experimental model, while contacts with lower acceptance frequency display high variability and false positive rates. We observed absence of contacts with high acceptance frequency for the IscS-frataxin pair (compared to the IscU-IscS case) (Fig S4A). The acceptance frequency, 68%, of the two most frequent contacts (Fig 5B) falls in the range where, in the case of IscU-IscS, most contacts are false positives. Therefore, even though the two contacts have geometrical compatibility, i.e. they could in principle be satisfied by a docked pose, their high statistical uncertainty prevents drawing conclusions about their biological relevance. In the case of interactions between frataxin and IscU, IPA identified three contacts with very high acceptance frequencies (>94%) (Fig 5C and Fig S4B) and potentially geometrically compatible with a docked complex. However, there is no overlap between these three coevolutionary predicted contacts and the interaction interface between frataxin and IscU in an available model of the IscU-IscS-frataxin trimer [37]. It must however be noted that the number of sequences in the IscU and IscS families are significantly higher than for the frataxin family. This will probably contribute to a higher statistical robustness of the predictions for the IscU-IscS complex.

## III. DISCUSSION

DCA is a powerful method, by now shown to be robust and reliable as long as a sufficiently high number of independent protein sequences are available [2,38,39]. In this work, we have interrogated evolution through DCA to gain new insights into the molecular machine involved in FeS cluster biogenesis. We selected three essential components: the scaffold protein IscU, the alternative scaffold IscA and the regulator of cluster formation, CyaY/frataxin. Besides the medical and biological interest of the latter, the choice of CyaY/frataxin revealed to be appropriate to validate the method for our purposes since this protein has a well compact and stable fold which presents a high structural conservation. Fewer CyaY/frataxin sequences are in agreement with the origin of this protein back only to the root of the alpha-beta-gamma proteobacteria, whereas, for instance, the IscU presence goes back to at least to the origin of bacteria. IscU is thus older of at least a couple of 100 million years (M. Huynen, personal communication). We nevertheless observed that, despite the relatively lower number of sequences, we can reproduce most features of the CyaY/frataxin fold, giving us confidence with the other two much better represented proteins. We then applied DCA to resolve questions which could allow us to understand cluster coordination and protein assembly of the other two proteins.

**FIG. 4.**
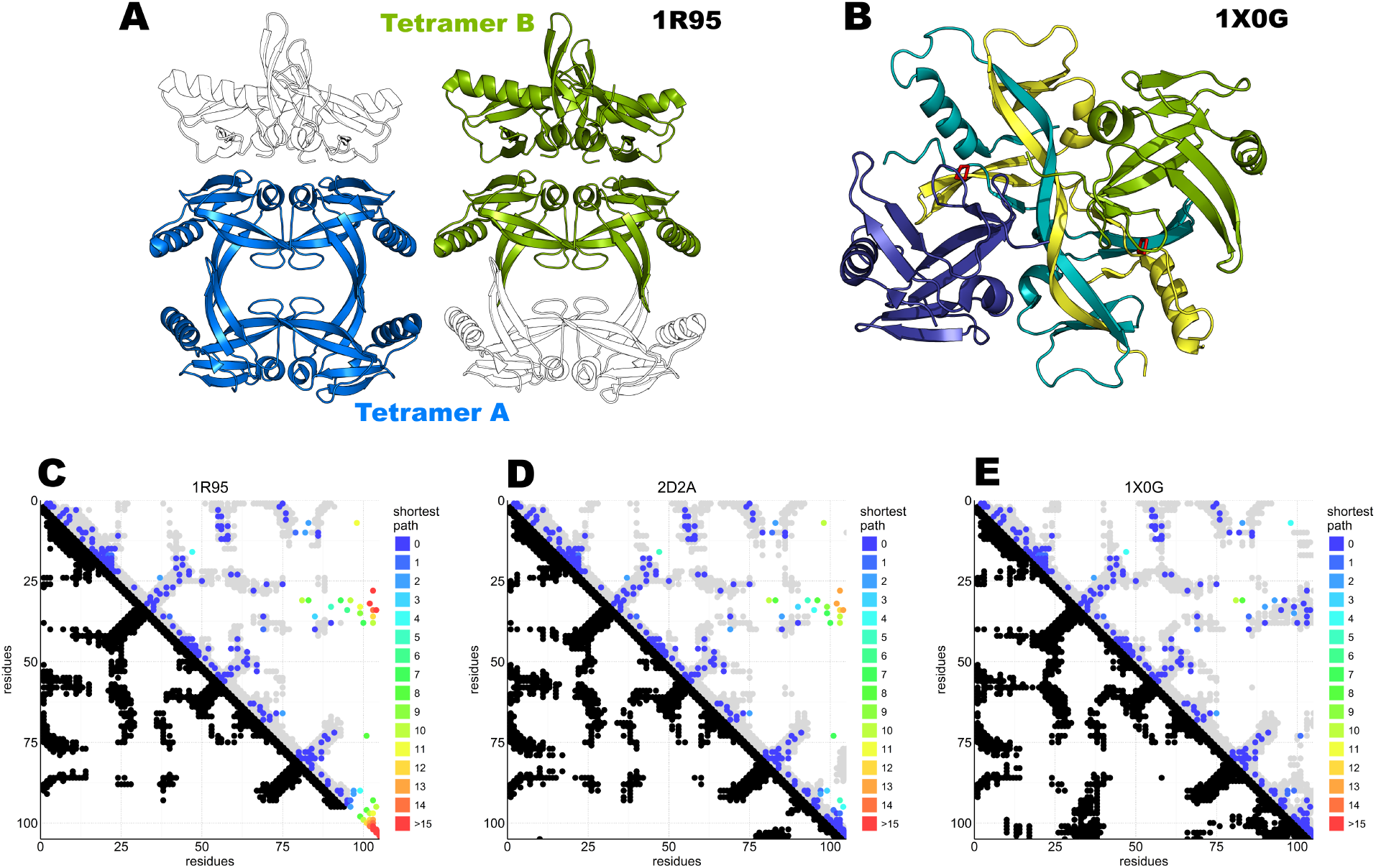
DCA analysis of IscA superimposed to available structures. A) IscA reference structure 1R95 with the two proposed tetramerization interfaces. The tetramer A (left) is the most broadly accepted biological unit. B) Domain swapped IscA tetramer (1X0G) bound to the FeS cluster. In shades of blue and green the two dimers. C) DCA predictions compared to the 1R95 reference structure. Most predictions are accurate, but the missing C-terminus hinders interpretation of the cluster binding site. DCA predictions compared to 1S98 are nearly identical and not shown. D) DCA predictions compared to the SufA 2D2A reference structure. Most predictions are accurate, but the model shows relevant differences in the C-terminus and for contacts between the terminal cysteine and the Cys35 regions. E) DCA predictions compared to 1X0G with domain-swapping. Nearly all predictions match the structure.

Much has been said about the presence of partially unstructured structures of IscU which could be in equilibrium with the fully folded form in solution [40]. There is no doubt that IscU is a marginally stable protein which, when in the absence of partners like zinc, the cluster or IscS is able to unfold not only at high but also at low temperatures [41]. The N-terminus is flexiblshe or in a conformational exchange in solution also in the presence of zinc. Nevertheless, we do not find traces of the unstructured conformation in our analysis, while the signal from the structured form is clear and unmistakable. Even more interestingly, we found for the first time some indication that directly supports experimentally the existence of a head-to-head IscU dimer whose interface would involve the conserved cysteines. This dimer was suggested to be the result of an oxidative event occurring at the later stages of FeS cluster formation, after the cluster-loaded IscU has detached from IscS [32]. This event would lead to the formation of a 4Fe4S cluster. IscU dimerization is the only way to reach sufficient coordination groups and enable formation of the 4Fe4S cubane which would instead be too unstable to be coordinated by the IscU monomer [41].

**FIG. 5.**
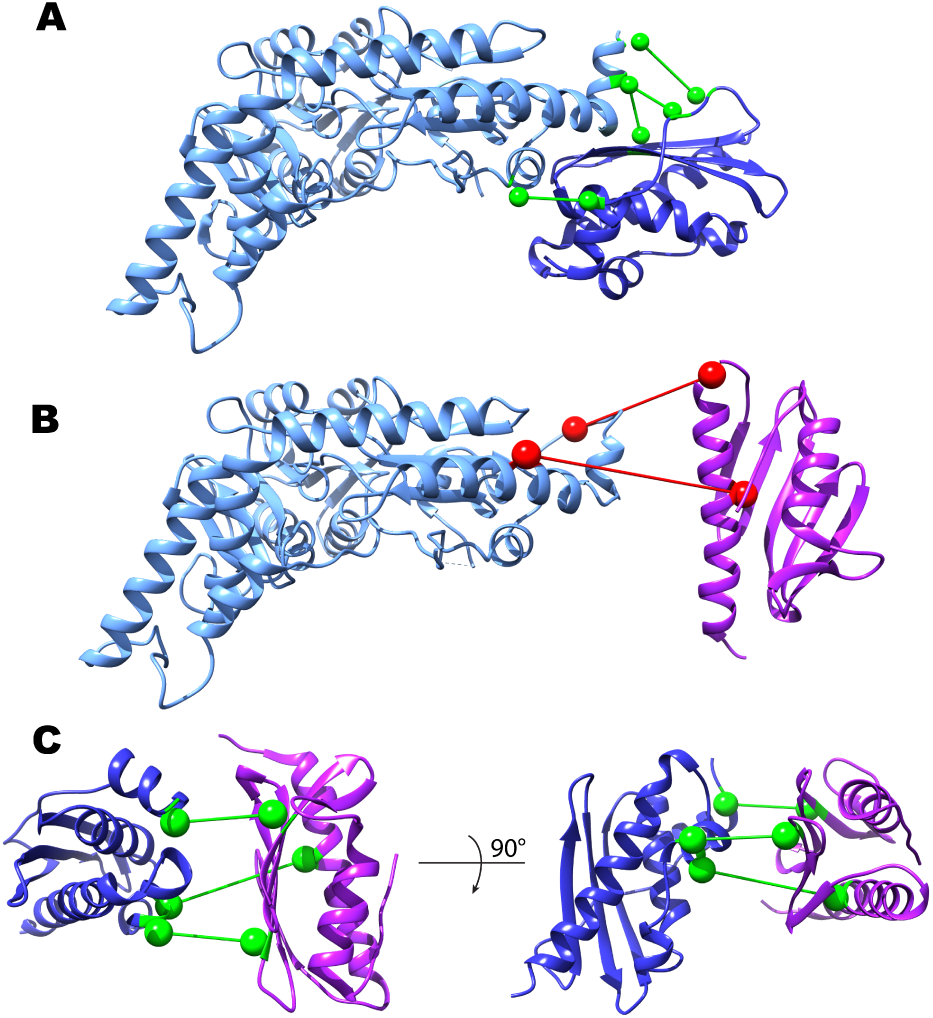
Inter-protein contact predictions for CyaY/frataxin, IscU and IscS. Inter-protein contacts predicted by IPA are shown in a ball-and-stick representation. The spheres are centered on the C*β*atoms (Cαfor glycine). Light Blue: IscS, Dark Blue: IscU, Purple: CyaY/frataxin. Contacts are colored according to their estimated robustness, based on the IscU-IscS reference case (Green: Robust contacts, acceptance frequency >85%; Red: Less robust contacts, acceptance frequency <85%). A) IscU-IscS interaction. The four contacts with highest acceptance frequency are shown. The IscU-IscS complex is drawn using the PDB 3LVL structure. B) Frataxin-IscS interaction. No robust contacts are predicted for the frataxin-IscS case. Reported are the two contacts with highest acceptance frequency (68%). C) Frataxin-IscU interaction. Three contacts have a high acceptance frequency (>94%).

DCA of IscA suggests new hypotheses on the structure of this otherwise still obscure protein. Because IscA binds both iron and FeS clusters, the protein has alternatively been suggested to be a scaffold protein or the carrier protein that delivers iron to the desulfurase [35,42,43]. What remains certain is that IscA contains three conserved cysteines, which are excellent candidates for both ion and cluster coordination. The crystal structures of IscA have been relatively uninformative both on the type of molecular assembly and on cluster/metal coordination. Our DCA data rely on a large number of sequences, just a little bit inferior to those retrieved for IscU. We observe a signal that is compatible with formation of the *αβ* fold observed in all available structures. However, we also observe contacts which cannot easily be explained by only one structure, suggesting the presence of several different species at least in the absence of cluster or cations. This is well consistent with our experimental evidence [44] which clearly supports the presence of an equilibrium between at least two species in a range of concentrations compatible with those expected in the cell. After analysing different structures we conclude that the co-presence of structures such like 1X0G and 1R95 would match what we observe in the DCA analysis. These conclusions strongly suggest that, while not necessarily giving domain swapping, we can envisage cluster coordination mediated by the dimeric form of IscA rather than the tetramer.

In conclusion, we found that DCA is a methodology which can enhance our knowledge on specific protein families and provide new information that can address unresolved questions. We can thus confidently add DCA to the tools which can allow us to study the FeS cluster machine.

### A. Materials and Methods

#### 1. Multiple Sequence Alignments

Multiple sequences alignments (MSAs) for each of the studied protein families were constructed using the following protocol: We first gathered all sequences from Uniprot with gene names corresponding to the canonical members of the families (CYAY or FXN for frataxin, ISCA for IscA, ISCS for IscS, ISCU for IscU). We then aligned the sequences in each seed using MAFFT (http://mat.cbrc.jp/alignment/software/) [45]. The resulting MSA was then used to generate a Hidden Markov Model using the HMMER package (http://hmmer.org/) [46] The Uniprot database was then searched using the HMMs to extract homologues sequences. The resulting MSAs were further filtered, removing all sequences containing more than 10% of gapped positions.

#### 2. Direct Coupling Analysis

DCA [2,3] was performed using an in-house code of the asymmetric version of the Pseudo-likelihood method to infer the parameters of the Potts Model [29,38]. Sequences were reweighted using a maximum 90% identity threshold. We used the L_2_ regularization parameters [38]. The DCA scores were taken as the Frobenius norm of the 20x20 coupling matrices J_ij_ of the Potts model (ignoring the couplings with the gap state) [47]. The average product correction term was subtracted [48]. The result was filtered to remove background and allow easier interpretation. The N most scoring predictions (N equals the MSA sequence length) were compared with the contact map of reference structures in which two residues were considered in contact if they have at least one atom 8.5 Åapart. Contacts between residues <5 amino-acids apart in the sequence were skipped to favor visualization of long range contact interactions.

#### 3. Iterative Paralog Matching and inter-protein predictions

To build matched MSAs of two interacting protein families (denoted A and B), we used the Iterative Paralog Matching (IPA) strategy [10]. The rationale of this procedure is to find the matching between paralogs of two protein families in an organism, such that the inter-protein coevolutionary signal is self-consistently maximized. The protocol can be summarized as follows: a random seed is built, such that for each organism, sequences of protein A are randomly matched with sequences of protein B. Mean-Field DCA [2] is used to infer the statistical model. The random seed is then discarded. The inferred couplings are then used to score all possible matchings of paralogs in all organisms. All potential matched sequence pairs are then ranked based on their inter-protein coevolution score, and a user defined number Ninc of the top ranking sequence pairs is added to the MSA, which will then be fed as input to MF-DCA for the next iteration. This procedure is repeated, increasing Ninc at each iteration, until the maximal number of sequences are matched. Finally, the best scoring MSA obtained by IPA is used as input to the Pseudo-Likelihood DCA method described above to perform contact prediction. This procedure is repeated N_*IPA*_ times, and for each realization, we recorded the inter-protein contacts with normalized DCA score above 0.8, an acceptance criterion introduced in [9]. We used N_*IPA*_=200 realizations for the IscU-IscS system, and N_*IPA*_=300 for the faster frataxin-IscU and frataxin-IscS systems. Finally, we ranked all inter-protein contacts by the normalized number of times they were accepted in the N_*IPA*_ realizations (acceptance frequency). Contacts being accepted more often across several IPA runs should reflect more robustness and higher statistical significance.

---

**FIG. S1.**
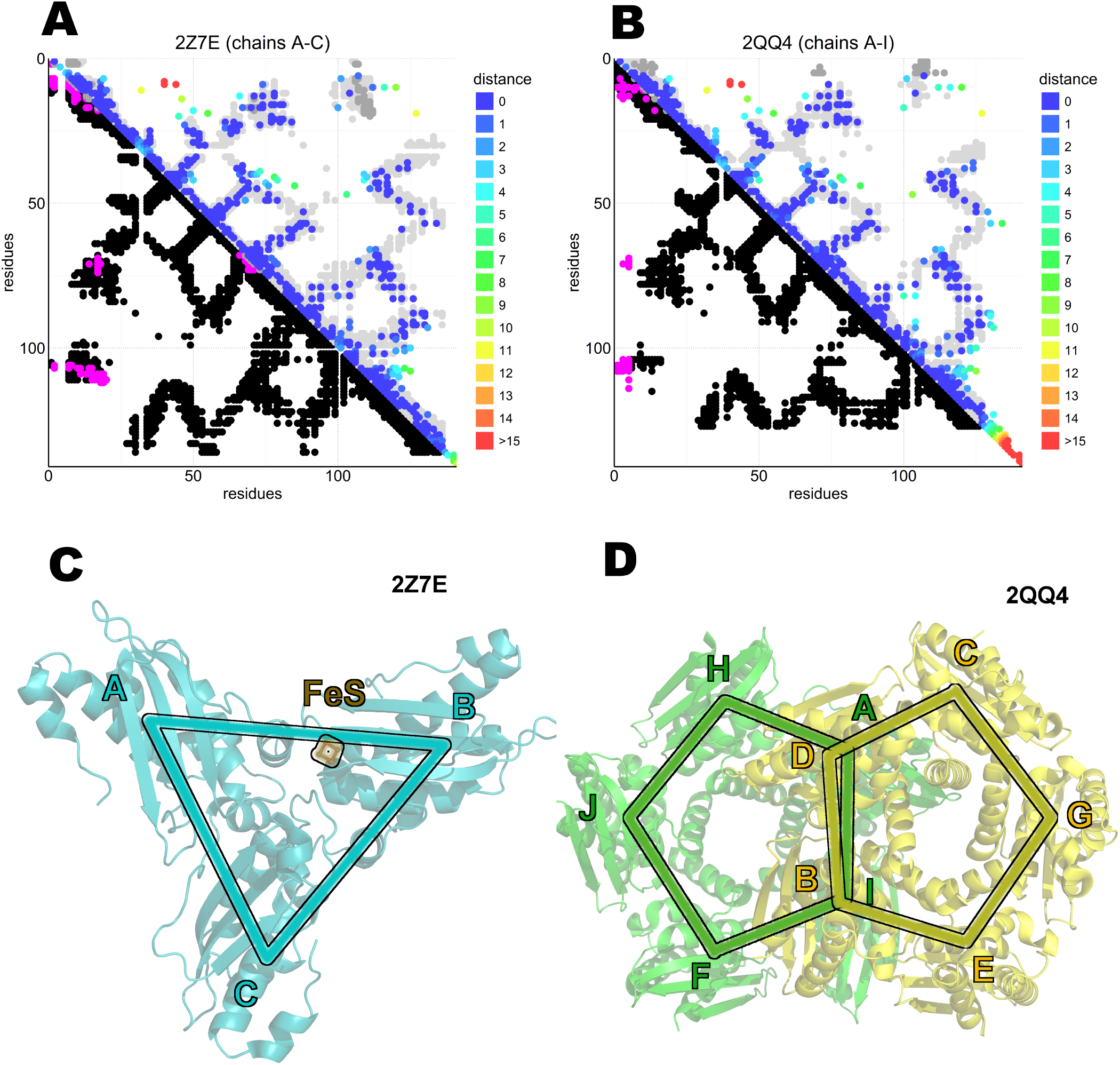
Interchain contacts of the multimeric IscU models. A-B) DCA on the IscU family over the available multimeric IscU structures. Each axis represent the position on the family consensus sequence from the N-to the C-terminus. In the bottom half of the plot, black coloured dot are placed where the coordinates define a pair of residues in contact in the same protomer of reference structures. In magenta, contacts between different protomers. In the top half of the plot the predicted DCA contacts are coloured by the shorted path from a reference matching position (gray dots on background. Gray dots are the same dots shown in the bottom half of the plots (black and magenta) but replotted on the top half to help the visualization of the predictions). A) 2Z7E with the interaction between A and C protomers. B) 2QQ4 with the interaction between A and I protomers. The other possible type of interaction surfaces (notably between chains A-D and B-I but also A-C and A-B) generated even less, unrelated, contacts (data not shown). C) Homotrimeric asymmetric iscU with FeS cluster (2Z7E). Only the B protomer is able to bind the FeS cluster. D) Homodecameric IscU (2QQ4).

**FIG. S2.**
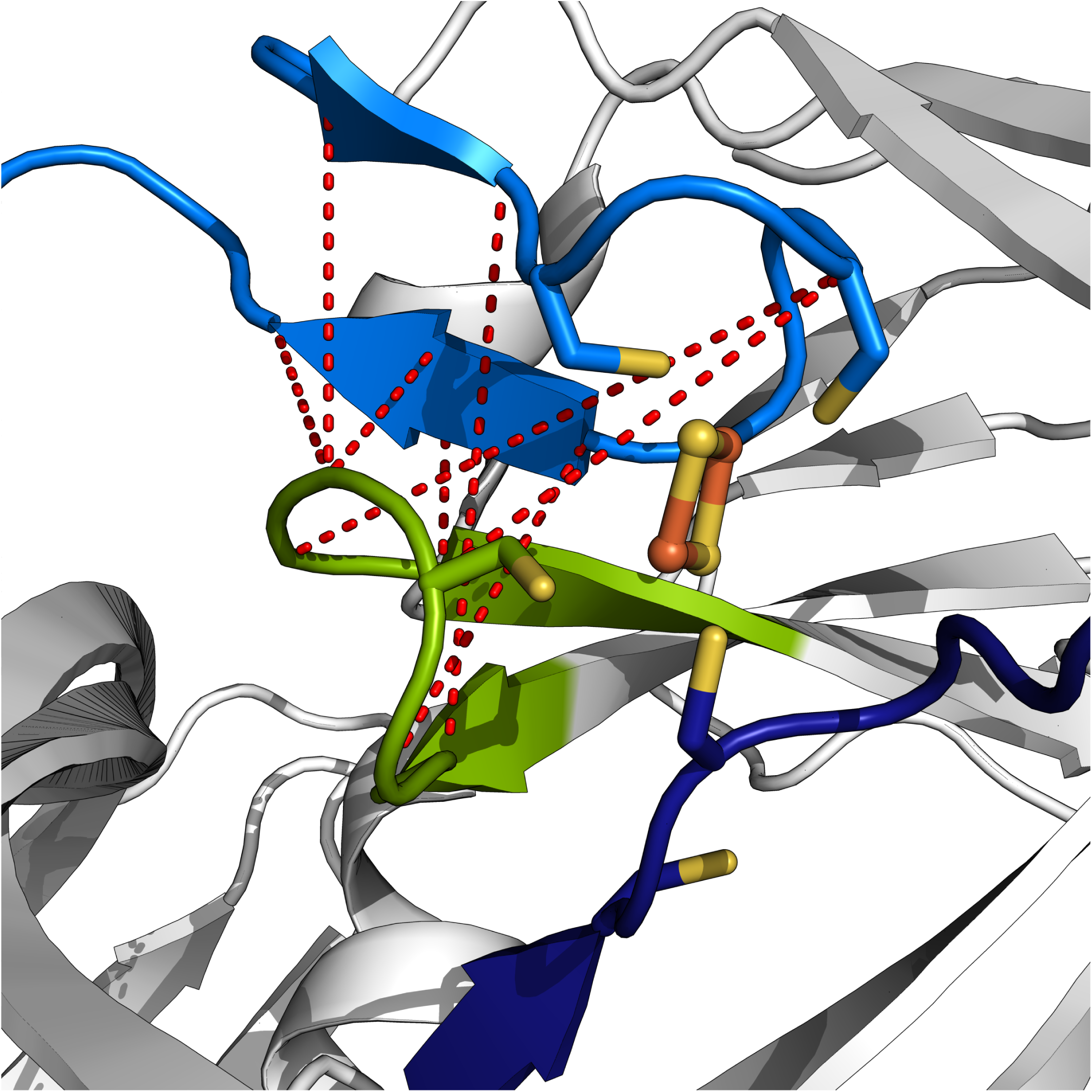
FeS cluster binding pocket of the asymmetrical IscA tetramer (1X0G).). 1X0G IscA tetramer with FeS cluster (in the middle, orange and yellow) bound to the cysteines of the C*x*_*n*_CGCG motif. In red are shown the DCA contacts between the loop containing the first cysteine of the motif (green) and the terminal segment containing the last two cysteines (cyan). A darker shade of blue dye the terminal segment on another protomer that contain the last cysteine bound to the FeS cluster. For simplicity, in the figure are displayed only the side-chains of the cysteines. The DCA predictions show several constrains that support the bending of the loop of the first cysteine toward the FeS binding site.

**FIG. S3.**
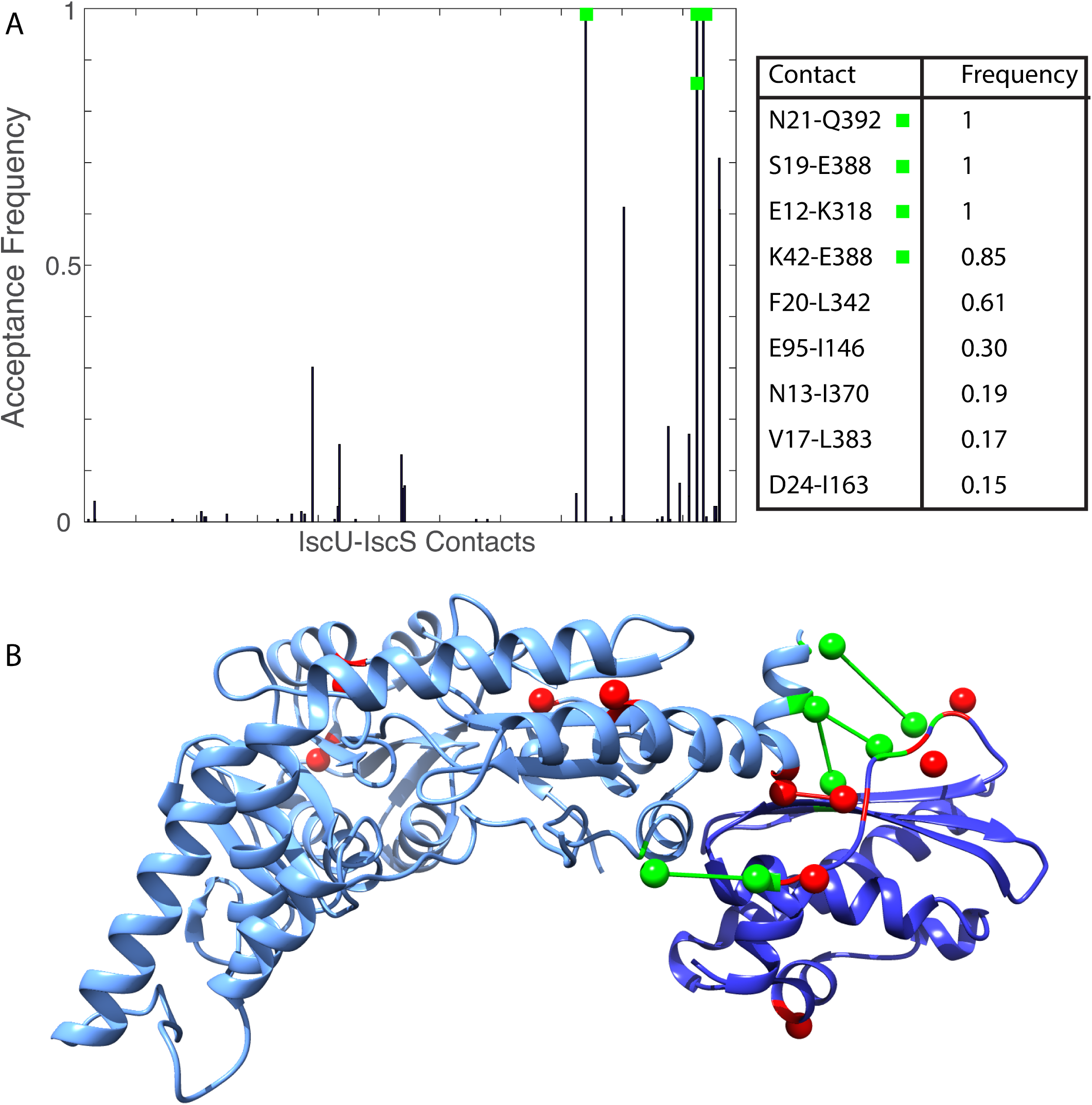
IPA calibration on the IscU-IscS complex. .A) (Left) Acceptance frequencies of the inter-protein contacts as estimated by N_*IPA*_=200 realizations of the IPA algorithm with random initial seed for the IscU-IscS system. The four contacts with highest acceptance frequency are true-positives in the reference structure of the complex (3LVL), and denoted with green squares. The horizontal axis is an arbitrary contact index. (Right) The list of the 9 predicted contacts with highest acceptance frequency. Note that two predicted involve positions with no mapping in the 3LVL structure and are not reporeted here. Both unmapped residues lie in the end of the C-terminal of IscS, in a particularly gapped region of the MSA. B) The 9 predicted IPA contacts, mapped on the PDB 3LVL structure. The spheres are centered on the C*β*atoms of the residue. The four highest contacts in A) are depicted in green, the remaining in red. Only contacts with C*β*-C*β* distance lower than 15 Å are depicted by sticks.

**FIG. S4.**
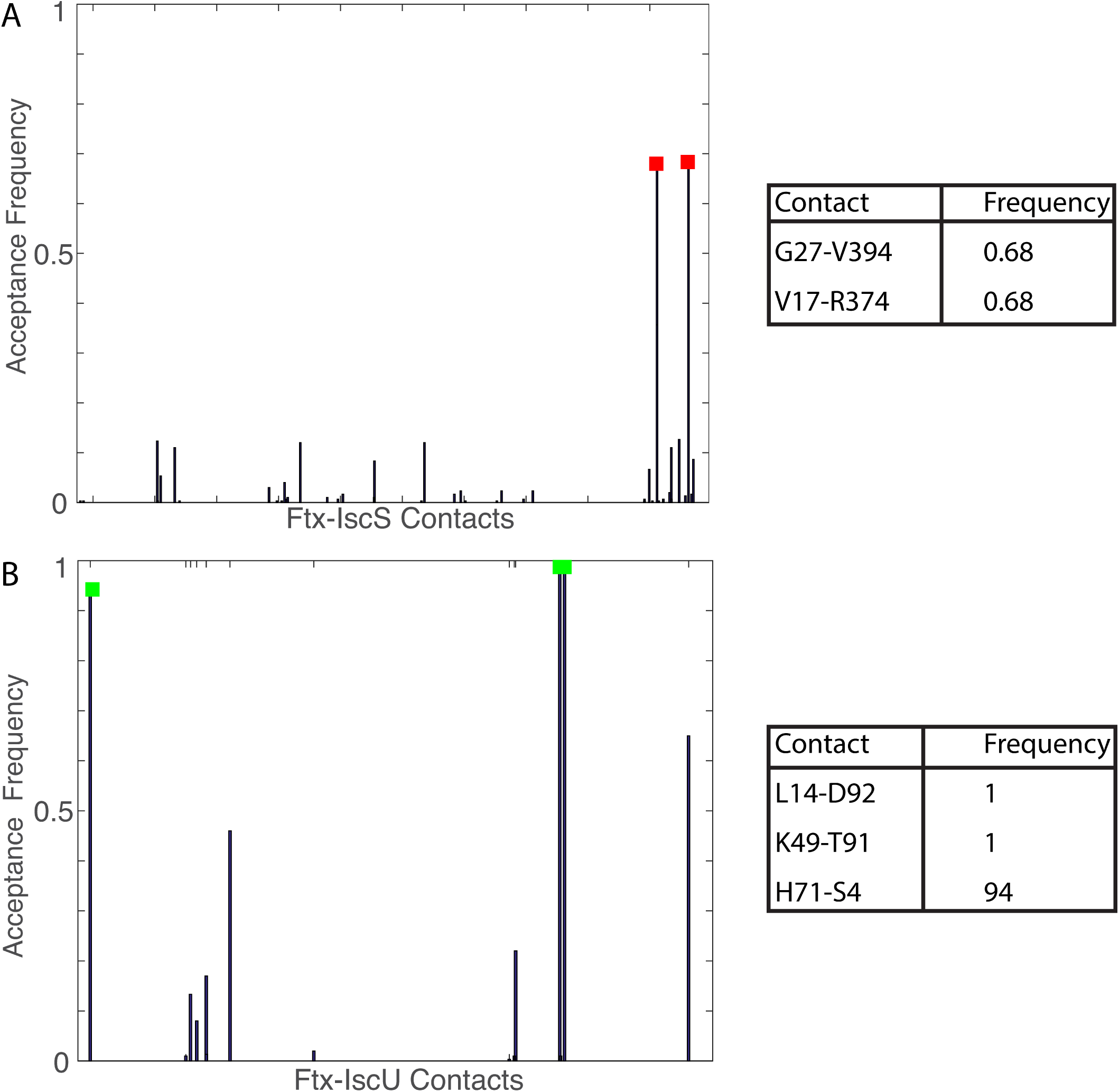
Selection Frequencies and contact list for Frataxin-IscU/IscS systems. A) (Left) Acceptance frequencies of the inter-protein contacts as estimated by N_*IPA*_=300 realizations of the IPA algorithm with random initial seed for the Frataxin-IscS system. The two contacts with highest acceptance frequency are highligted by red squares. The horizontal axis is an arbitrary contact index. (Right) The list of the two inter-protein contacts with highest acceptance frequency. B) Acceptance frequencies of the inter-protein contacts as estimated by N_*IPA*_=300 realizations of the IPA algorithm with random initial seed for the Frataxin-IscU system. The three contacts with highest acceptance frequency are highligted by green squares. The horizontal axis is an arbitrary contact index. (Right) The list of the three inter-protein contacts with highest acceptance frequency.

## REFERENCES

1. GöbelU, Sander C, Schneider R, Valencia A. Correlated mutations and residue contacts in proteins. Proteins Struct Funct Genet. 1994;18: 309–317. doi:10.1002/prot.340180402

2. Morcos F, Pagnani A, Lunt B, Bertolino A, Marks DS, Sander C, et al. Direct-coupling analysis of residue coevolution captures native contacts across many protein families. Proc Natl Acad Sci U S A. 2011;108: E1293–301. doi:10.1073/pnas.1111471108

3. Weigt M, White R a, Szurmant H, Hoch J a, Hwa T. Identification of direct residue contacts in protein-protein interaction by message passing. Proc Natl Acad Sci U S A. 2009;106: 67–72. doi:10.1073/pnas.0805923106

4. Malinverni D, Marsili S, Barducci A, De Los Rios P. Large-Scale Conformational Transitions and Dimerization Are Encoded in the Amino-Acid Sequences of Hsp70 Chaperones. PLoS Comput Biol. 2015;11: 1–15. doi:10.1371/journal.pcbi.1004262

5. Hopf TA, Morinaga S, Ihara S, Touhara K, Marks DS, Benton R. Amino acid coevolution reveals three-dimensional structure and functional domains of insect odorant receptors. Nat Commun. 2015; 6:6077. doi:10.1038/ncomms7077

6. Espada R, Parra RG, Mora T, Walczak AM, Ferreiro DU. Capturing coevolutionary signals inrepeat proteins. BMC Bioinformatics. 2015;16: 207. doi:10.1186/s12859-015-0648-3

7. Sutto L, Marsili S, Valencia A, Gervasio FL. From residue co-evolution to protein conformational ensembles and functional dynamics. Proc Natl Acad Sci U S A. 2015;112: 13567–72. doi:10.1073/pnas.1508584112

8. Ovchinnikov S, Kamisetty H, Baker D. Robust and accurate prediction of residue-residue interactions across protein interfaces using evolutionary information. Elife. 2014;2014: 1–21. doi:10.7554/eLife.02030

9. Hopf TA, Schrfe CPI, Rodrigues JPGLM, Green AG, Kohlbacher O, Sander C, et al. Sequence co-evolution gives 3D contacts and structures of protein complexes. Elife. 2014;3: e03430. doi:10.7554/eLife.03430

10. Bitbol A-F, Dwyer RS, Colwell LJ, Wingreen NS. Inferring interaction partners from protein sequences. Proc Natl Acad Sci U S A. National Academy of Sciences; 2016;113: 12180–12185. doi:10.1073/pnas.1606762113

11. Procaccini A, Lunt B, Szurmant H, Hwa T, Weigt M. Dissecting the specificity of protein-protein interaction in bacterial two-component signaling: Orphans and crosstalks. PLoS One. 2011;6. doi:10.1371/journal.pone.0019729

12. Beilschmidt LK, Puccio HM. Mammalian Fe-S cluster biogenesis and its implication in disease. Biochimie. 2014. pp. 48–50. doi:10.1016/j.biochi.2014.01.009

13. Rouault TA. Mammalian iron-sulphur proteins: novel insights into biogenesis and function. Nat Rev Mol Cell Biol. Nature Publishing Group; 2015;16: 45–55. doi:10.1038/nrm3909

14. Bothe JR, Tonelli M, Ali IK, Dai Z, Frederick RO, Westler WM, et al. The Complex Energy Landscape of the Protein IscU. Biophys J. The Authors; 2015;109: 1019–1025. doi:10.1016/j.bpj.2015.07.045

15. Gibson TJ, Koonin E V., Musco G, Pastore A, Bork P. Friedreich’s ataxia protein: Phylogenetic evidence for mitochondrial dysfunction. Trends Neurosci. 1996;19: 465–468. doi:10.1016/S0166-2236(96)20054-2

16. Adinol S, Iannuzzi C, Prischi F, Pastore C, Iametti S, Martin SR, et al. Bacterial frataxin CyaY is the gatekeeper of iron-sulfur cluster formation catalyzed by IscS. Nat Struct Mol Biol. 2009;16: 390–396. doi:10.1038/nsmb.1579

17. Shi R, Proteau A, Villarroya M, Moukadiri I, Zhang L, Trempe JF, et al. Structural basis for Fe-S cluster assembly and tRNA thiolation mediated by IscS protein-protein interactions. PLoS Biol. 2010;8. doi:10.1371/journal.pbio.1000354

18. Prischi F, Konarev P V, Iannuzzi C, Pastore C, Adinol S, Martin SR, et al. Structural bases for the interaction of frataxin with the central components of iron-sulphur cluster assembly. Nat Commun. Nature Publishing Group; 2010;1: 95. doi:10.1038/ncomms1097

19. Iannuzzi C, Adinolfi S, Howes BD, Garcia-Serres R, Clémancey M, Latour JM, et al. The role of cyay in iron sulfur cluster assembly on the e. coli iscu scaffold protein. PLoS One. 2011;6. doi:10.1371/journal.pone.0021992

20. Tsai CL, Barondeau DP. Human frataxin is an allosteric switch that activates the Fe-S cluster biosynthetic complex. Biochemistry. 2010;49: 9132–9139. doi:10.1021/bi1013062

21. Gakh O, Bedekovics T, Duncan SF, Smith IV DY, Berkholz DS, Isaya G. Normal and Friedreich ataxia cells express different isoforms of frataxin with complementary roles in iron-sulfur cluster assembly. J Biol Chem. 2010;285: 38486–38501. doi:10.1074/jbc.M110.145144

22. Kaut A, Lange H, Diekert K, Kispal G, Lill R. Isa1p is a component of the mitochondrial machinery for maturation of cellular iron-sulfur proteins and requires conserved cysteine residues for function. J Biol Chem. 2000;275: 15955–15961. doi:10.1074/jbc.M909502199

23. Prischi F, Giannini C, Adinol S, Pastore A. The N-terminus of mature human frataxin is intrinsically unfolded. FEBS J. 2009;276: 6669–6676. doi:10.1111/j.1742-4658.2009.07381.x

24. Popovic M, Sanfelice D, Pastore C, Prischi F, Temussi PA, Pastore A. Selective observation of the disordered import signal of a globular protein by in-cell NMR: The example of frataxins. Protein Sci. 2015;24: 996–1003. doi:10.1002/pro.2679

25. Adinol S, Nair M, Politou A, Bayer E, Martin S, Temussi P, et al. The factors governing the thermal stability of frataxin orthologues: How to increase a protein’s stability. Biochemistry. 2004;43: 6511–6518. doi:10.1021/bi036049

26. Pastore C, Franzese M, Sica F, Temussi P, Pastore A. Understanding the binding properties of an unusual metal-binding protein - A study of bacterial frataxin. FEBS J. 2007;274: 4199–4210. doi:10.1111/j.1742-4658.2007.05946.x

27. Nair M, Adinol S, Pastore C, Kelly G, Temussi P, Pastore A. Solution structure of the bacterial frataxin ortholog, CyaY: Mapping the iron binding sites. Structure. 2004;12: 2037–2048. doi:10.1016/j.str.2004.08.012

28. Huynen MA, Snel B, Bork P, Gibson TJ. The phylogenetic distribution of frataxin indicates a role in iron-sulfur cluster protein assembly. Hum Mol Genet. 2001;10: 2463–8. doi:10.1093/hmg/10.21.2463

29. Balakrishnan S, Kamisetty H, Carbonell JG, Lee S-I, Langmead CJ. Learning generative models for protein fold families. Proteins. 2011;79: 1061–78. doi:10.1002/prot.22934

30. Liu J, Oganesyan N, Shin DH, Jancarik J, Yokota H, Kim R, et al. Structural characterization of an iron-sulfur cluster assembly protein IscU in a zinc-bound form. Proteins Struct Funct Genet. 2005;59: 875–881. doi:10.1002/prot.20421

31. Adrover M, Howes BD, Iannuzzi C, Smulevich G, Pastore A. Anatomy of an iron-sulfur cluster scaffold protein: Understanding the determinants of [2Fe-2S] cluster stability on IscU. Biochim Biophys Acta - MolCell Res. 2015;1853: 1448–1456. doi:10.1016/j.bbamcr.2014.10.023

32. Chandramouli K, Unciuleac MC, Naik S, Dean DR, Boi HH, Johnson MK. Formation and properties of [4Fe-4S] clusters on the IscU scaffold protein. Biochemistry. 2007;46: 6804–6811. doi:10.1021/bi6026659

33. Bilder PW, Ding H, Newcomer ME. Crystal structure of the ancient, Fe-S scaffold IscA reveals a novel protein fold. Biochemistry. 2004;43: 133–9. doi:10.1021/bi035440s

34. Cupp-Vickery JR, Silberg JJ, Ta DT, Vickery LE. Crystal structure of IscA, an iron-sulfur cluster assembly protein from Escherichia coli. J Mol Biol. 2004;338: 127–137. doi:10.1016/j.jmb.2004.02.027

35. Krebs C, Agar JN, Smith AD, Frazzon J, Dean DR, Huynh BH, et al. IscA, an alternate scaffold for Fe-S cluster biosynthesis. Biochemistry. 2001;40: 14069–14080. doi:10.1021/bi015656z

36. Gueudré T, Baldassi C, Zamparo M, Weigt M, Pagnani A. Simultaneous identification of specifically interacting paralogs and interprotein contacts by direct coupling analysis. Proc Natl Acad Sci. 2016;113: 12186–12191. doi:10.1073/pnas.1607570113

37. di Maio D, Chandramouli B, Yan R, Brancato G, Pastore A. Understanding the role of dynamics in the iron sulfur cluster molecular machine. Biochim Biophys Acta - Gen Subj. The Authors; 2016; doi:10.1016/j.bbagen.2016.07.020

38. Ekeberg M, Lvkvist C, Lan Y, Weigt M, Aurell E. Improved contact prediction in proteins: Using pseudolikelihoods to infer Potts models. Phys Rev E - Stat Nonlinear, Soft Matter Phys. 2013;87. doi:10.1103/PhysRevE.87.012707

39. Marks DS, Hopf T a, Sander C. Protein structure prediction from sequence variation. Nat Biotechnol. 2012;30: 1072–80. doi:10.1038/nbt.2419

40. Markley JL, Kim J, Dai Z, Bothe JR, Cai K, Frederick RO, et al. Metamorphic protein IscU alternates conformations in the course of its role as the scaffold protein for iron-sulfur cluster biosynthesis and delivery. FEBS Lett. Federation of European Biochemical Societies; 2013;587: 1172–1179. doi:10.1016/j.febslet.2013.01.003

41. Iannuzzi C, Adrover M, Puglisi R, Yan R, Temussi PA, Pastore A. The role of zinc in the stability of the marginally stable IscU scaffold protein. Protein Sci. 2014;23: 1208–1219. doi:10.1002/pro.2501

42. Ollagnier-De-Choudens S, Mattioli T, Takahashi Y, Fontecave M. Iron-sulfur cluster assembly. Characterization of IscA and evidence for a specific and functional complex with ferredoxin. J Biol Chem. 2001;276: 22604–22607. doi:10.1074/jbc.M102902200

43. Ding H, Yang J, Coleman LC, Yeung S. Distinct iron binding property of two putative iron donors for the iron-sulfur cluster assembly: IscA and the bacterial frataxin ortholog CyaY under physiological and oxidative stress conditions. J Biol Chem. 2007;282: 7997–8004. doi:10.1074/jbc.M609665200

44. Popovic M, Pastore A. Chemical shift assignment of the alternative scaffold protein IscA. Biomol NMR Assign. Springer Netherlands; 2016;10: 227–231. doi:10.1007/s12104-016-9672-0

45. Katoh K. MAFFT: a novel method for rapid multiple sequence alignment based on fast Fourier transform. Nucleic Acids Res. 2002;30: 3059–3066. doi:10.1093/nar/gkf436

46. Finn RD, Clements J, Eddy SR. HMMER web server: Interactive sequence similarity searching. Nucleic Acids Res. 2011;39. doi:10.1093/nar/gkr367

47. Feinauer C, Skwark MJ, Pagnani A, Aurell E. Improving Contact Prediction along Three Dimensions. PLoS Comput Biol. 2014;10. doi:10.1371/journal.pcbi.1003847

48. Dunn SD, Wahl LM, Gloor GB. Mutual information without the influence of phylogeny or entropy dramatically improves residue contact prediction. Bioinformatics. 2008;24: 333–340. doi:10.1093/bioinformatics/btm604

